# Intranasal oxytocin enhances social preference for parents over peers in male but not female peri-adolescent California mice (*Peromyscus californicus*)

**DOI:** 10.1101/2022.09.12.507587

**Authors:** Caleigh D. Guoynes, Catherine A. Marler

## Abstract

Peri-adolescence is a critical developmental stage marked by profound changes in the valence of social interactions with parents and peers. We hypothesized that the oxytocin (OXT) and vasopressin (AVP) systems, known for influencing social behavior, would be involved in the maintenance and breaking of bonding behavior expressed by peri-adolescent males and females. In rodents, OXT is associated with mother-pup bonding and may promote social attachment to members of the natal territory. AVP, on the other hand, can act in contrasting ways to OXT and has been associated with aggression and territoriality. Specifically, we predicted that in peri-adolescent male and female juveniles of the biparental and territorial California mouse (*Peromyscus californicus*), a) OXT would increase the social preferences for the parents over unfamiliar age-matched peers (one male and one female), and b) AVP would break the parent-offspring bond and either increase time in the neutral chamber and/or approach to their unfamiliar and novel peers. We examined anxiety and exploratory behavior using an elevated plus maze and a novel object task as a control. Peri-adolescent mice were administered an acute intranasal (IN) treatment of 0.5 IU/kg IN AVP, 0.5 IU/kg IN OXT, or saline control; five minutes later, the behavioral tests were conducted. As predicted, we found that IN OXT enhanced social preference for parents; however, this was only in male and not female peri-adolescent mice. IN AVP did not influence social preference in either sex. These effects appear specific to social behavior and not anxiety, as neither IN OXT nor AVP influenced behavior during the elevated plus maze or novel object tasks. To our knowledge, this is the first evidence indicating that OXT may play a role in promoting peri-adolescent social preferences for parents and delaying weaning in males.

**HIGHLIGHTS:**

- In a 3-chambered choice test, peri-adolescent female and male California mice prefer their parents over peers or an empty chamber
- Intranasal oxytocin (IN OXT) enhances male but not female peri-adolescent social preference for their parents
- Intranasal arginine vasopressin (IN AVP) did not influence social preference in either sex
- Neither IN OXT nor AVP alter peri-adolescent behavior in an elevated plus maze or novel object task
- OXT may play a role in delaying weaning in males

## INTRODUCTION

Across a wide variety of species, the developmental stage of adolescence is highly conserved (Spear, 2007). Mammals with shorter lifespans and less complex social environments enter puberty more quickly (Brenhouse & Andersen, 2011). For these species, the cost of prolonging time in their natal territory and competing with their siblings and parents for resources outweighs the benefits of parental food resources, protection, and social learning. However, a longer buffer period between puberty and adulthood may be advantageous for some species. For example, in species with dominance hierarchies, prolonged adolescence likely helps to attain higher social ranks and greater mating opportunities (Steinberg, 2010; Duell & Steinberg, 2019). Resource availability and group social composition may change across births from the same parent or across generations as areas experience drought or other cyclic environmental pressures (Prugh et al., 2018). Plasticity in weaning time could afford adolescents more time to prepare to explore social challenges in and out of their natal territory, migrate, and establish new territories when conditions are less favorable.

While environmental pressures guide a peri-adolescent mammal’s weaning time, their social bond with their parent(s) may also be a proximate mechanism influencing weaning time. Among several candidate neurochemicals, there are two groups of likely hormonal mechanisms that could affect this process: one is the sex steroid hormones that influence maturation and the process of puberty (Romeo, 2003; Forbes & Dahl, 2010; Delevich et al., 2021) and the second is oxytocin (OXT) that influences bonding (Carter, 1992; Kendrick, 2000; Liu & Wang, 2003; Nagasawa et al., 2012; Loth & Donaldson, 2021), but is also influenced by those sex steroid hormones (Shapiro et al., 2000; Murakami et al., 2011). Dramatic steroid hormone changes are associated with the transition from being a juvenile to becoming an adult (Peper et al., 2011; Koolschijn et al., 2014; Trova et al., 2021), and these hormones can also drive the regulation of OXT and a similar neuropeptide, vasopressin (AVP) (DeVries et al., 1994). OXT and AVP influence social behavior and preference across a wide variety of mammalian species, including rodents (Kent et al., 2013; Lukas & Neumann, 2014; Wood et al., 2014; Duque-Wilckens et al., 2018; Williams et al., 2020; Bester-Meredith & Marler, 2001; Guoynes et al., 2018; Lukas et al., 2011; Albers, 2012; Winslow et al., 1993; Wang et al., 1999; Keverne & Curley, 2004; Prounis et al., 2018; Albers & Bamshed, 1999; Taylor et al., 2022; Taylor et al., 2017), and nonhuman and human primates (Taylor & French, 2015; Jarcho et al., 2011; Baxter et al., 2020; Crockford et al., 2013; Staes et al., 2015; Heinrichs et al., 2009; Skuse et al., 2014; Maud et al., 2018; Bredewold & Veneema, 2018; Zhuang et al., 2021). Here we focus on the OXT and AVP systems as neurochemical candidates for influencing weaning time in mammals. In mothers, the physiological release of oxytocin (OXT) promotes milk letdown (Nishimori et al., 1996; Young et al., 1997), increases circulating OXT via increased pup suckling (Febo et al., 2005; Neumann et al., 1993), and changes OXTR receptor expression in social brains areas such as the nucleus accumbens, ventral tegmental area, bed stria of the nucleus terminalis, ventromedial hypothalamus, lateral septum (Keebaugh et al., 2015; Insel, 1990; Pedersen et al., 1994; Curley et al., 2012). In the early stages of rearing, a single intranasal dose of OXT is enough to promote increased maternal communication and care in California mice (Guoynes & Marler, 2021). These studies suggest a link between the OXT-driven positive feedback mechanisms associated with milk let-down/suckling and the mother-offspring bond.

Despite the above research on the effect of OXT on bond initiation in mothers, the neurobiological mechanisms that maintain the parent-offspring bond through peri-adolescence are not well understood. Is the bond maintained by the parent(s), the offspring, or both? Here we initially address this question through a mechanistic approach; we ask whether OXT might induce flexibility in weaning time. In young female and male rats, post-weaning isolation decreases OXT receptor expression in the nucleus accumbens and increases aggression, suggesting that higher OXT receptivity in the nucleus accumbens may be important for prosocial behavior (de Moura Oliveira et al., 2019). When offspring transition from the juvenile period to adulthood, OXTR expression decreases in the lateral septum, an area known for its role in integrating sensory information and prior social experiences (Smith et al., 2017). Therefore, it is possible that alterations to the OXT system not only facilitate changes to the mother’s social preference and behavior but also in her offspring’s preferences and behavior. Although our focus is on OXT and AVP, we also acknowledge the close connections between the sex steroid hormones and the OXT system (McCarthy et al., 1996; Frayne & Nicholson, 1995; Young et al., 1998; Murakami et al., 2011; Acevedo-Rodriguez et al., 2015) that underscore the possibility that the oxytocin system will affect females and males differently.

To test the hypothesis that OXT influences offspring-parent attachment during peri-adolescence, we use California mice (*Peromyscus californicus*), a monogamous, biparental rodent species whose social behavior can be modulated by OXT manipulations (Duque-Wilckens et al., 2018; Perea-Rodriguez et al., 2015; Guoynes & Marler, 2021; Monari et al., 2021; Guoynes & Marler, 2022; Gubernick et al., 1995; Williams et al., 2020; Steinman et al., 2019; Yohn et al., 2018). Like other species, the young mature and migrate from their natal territory. There is no published evidence of overlapping generations, and females and males disperse from the nest (Ribble, 1992). Interestingly, females disperse farther from their natal territory than males (Ribble, 1992), suggesting male juveniles may be less inclined to explore and show greater parent-offspring proximity and attachment behavior. In addition to testing the effect of OXT on parent-offspring bonding, we also wanted to examine whether AVP influenced parent-offspring bonding due to its role in territoriality and aggression in California mice (Bester-Meredith & Marler, 2001; Frazier et al., 2006; Yohn et al., 2017). We predicted that while OXT would maintain the parent-offspring bond, AVP may erode the parent-offspring bond by increasing territoriality in the peri-adolescents. Moreover, we expected that if any sex showed a stronger or more flexible bond with the parents, it would be males because of staying closer to the natal territory. We tested these predictions using a three-chambered choice test in which peri-adolescents, close to weaning age, chose between spending time with two sets of social partners: their parents versus an unrelated, age-matched novel female and male. Furthermore, to test whether IN OXT and AVP are specific to peri-adolescent social behavior, we also tested the effect of IN OXT and IN AVP on two nonsocial tasks that assess anxiety and exploration: the elevated plus maze and the novel object task. We predicted that since these tasks do not involve social stimuli, we would not see an effect of either IN OXT or AVP.

This study is the first to examine how OXT and AVP influence the maintenance of offspring-parent social preferences during peri-adolescence. This work adds to the literature on the mechanisms underlying social bonds within family units, such as pair bonds and maternal and paternal bonds to their offspring.

## METHODS

### Animals

For our focal animals, we used 99 female and male *P. californicus* aged postnatal day (PND) 24-26 in litters 5-16 from 17 different breeding pairs. PND 24-26 represents early weaning and, therefore, will be referred to as peri-adolescence in California mice because they are typically weaned on PND 28-30. In contrast, other small rodents used in social behavior research, such as house mice (*Mus musculus*) and prairie voles (*Microtus ochrogaster*), are both typically weaned around PND 21 (Bechard & Mason, 2010; Guoynes et al., 2018; Horii-Hayashi et al., 2013). These animals were divided into two groups, with 44 animals in Group 1 that were randomly assigned to three treatment groups (OXT, AVP, and CTRL) and 55 animals in Group 2 that were randomly assigned to three treatment groups (OXT, AVP, and CTRL). For stimulus animals in the Group 1 study, we used 48 peri-adolescent stimulus animals aged PND 24-31 and 17 sets of parents (34 mice) aged 10-24 months that were unrelated by two generations. Parents were reused across the study, but never more than five times as stimulus animals, and offspring from one set of parents were never assigned to the same treatment group. One to 11 pups were used from each breeder pair, with a mean of 5.94 and a standard deviation of 2.51 pups used per breeder pair. Before testing, peri-adolescent California mice were housed in breeding cages with their parents and 2-4 siblings (48 × 27 × 16 cm) under a 14L: 10D light cycle with lights off at 1:00 pm. For testing, peri-adolescent mice were randomly assigned to Group 1 and Group 2, with Group 1 exposed to a parent-peer preference test and Group 2 exposed to an elevated plus maze test and novel object test. Individual mice were then randomly assigned to the following treatments (described in detail below) with the stated sample sizes. In Group 1, sample sizes were: CTRL female (N=8), OXT female (N=8), AVP female (N=7), CTRL male (N=7), OXT male (N=8), and AVP male (N=6). In Group 2 sample sizes were: CTRL female (N=10), OXT female (N=10), AVP female (N=9), CTRL male (N=8), OXT male (N=9), AVP male (N=9). Animals were maintained per the National Institute of Health Guide for the Care and Use of Laboratory Animals and the University of Wisconsin-Madison Institutional Animal Care and Use Committee approved this research.

### Intranasal Oxytocin (OXT) and Vasopressin (AVP) Preparation

Female and male mice were infused intranasally with 17.5 uL of either sterile saline, OXT (0.5 IU/kg), or AVP (0.5 IU/kg) (Bachem, Torrance, California) (as used in Guoynes & Marler, 2021, 2022; Monari et al., 2021). The 0.5 IU/kg dose is relatively low and acts like an acute pulse or bolus release of neuropeptide (Guoynes & Marler, 2021). This dose is also analogous to those used in human clinical trials (Anagnostou et al., 2014; Huang et al., 2021). OXT and AVP were dissolved in saline, prepared in two separate large batches, aliquoted into small plastic tubes, and frozen at 20°C. Treatments, including saline control, were defrosted just before administration. A blunt cannula needle (33-gauge, 2.8 mm length; Plastics One, Roanoke, Virginia) was attached to cannula tubing, flushed, filled with the compound, then attached to an airtight Hamilton syringe (Bachem, Torrance, California). The animal was scruffed, and 17.5 uL of the compound was expelled dropwise through the cannula needle and allowed to absorb into the nasal mucosa (∼10-20 seconds). One person conducted all IN administrations throughout the study to maintain consistency in handling and IN infusion. We chose to use IN delivery method because it has been shown to reach the brain in several species (Lee et al., 2020; Smith et al., 2019; Neumann et al., 2013; Striepens et al., 2013; Lee et al., 2018; Oppong-Damoah et al., 2019; Freeman et al., 2016) and is less invasive than intracranial injections (see Guoynes & Marler, 2021 and Guoynes & Marler, 2022).

### Behavioral Tests

To assess peri-adolescent social preference and exploration, we assigned peri-adolescent mice to one of two groups. Regardless of group assignment, each mouse was weaned from its home cage 24 hours prior to testing (PND 23-25) and singly housed in a new home cage (30 × 19 × 13 cm). In Group 1, mice were tested in a 30-min parent-peer preference test as described below. In Group 2, mice were tested in the elevated plus maze test and then immediately tested in the novel object test for a cumulative total of 15 min of behavioral testing as described below **(Fig. 1**).

**Figure 1.**
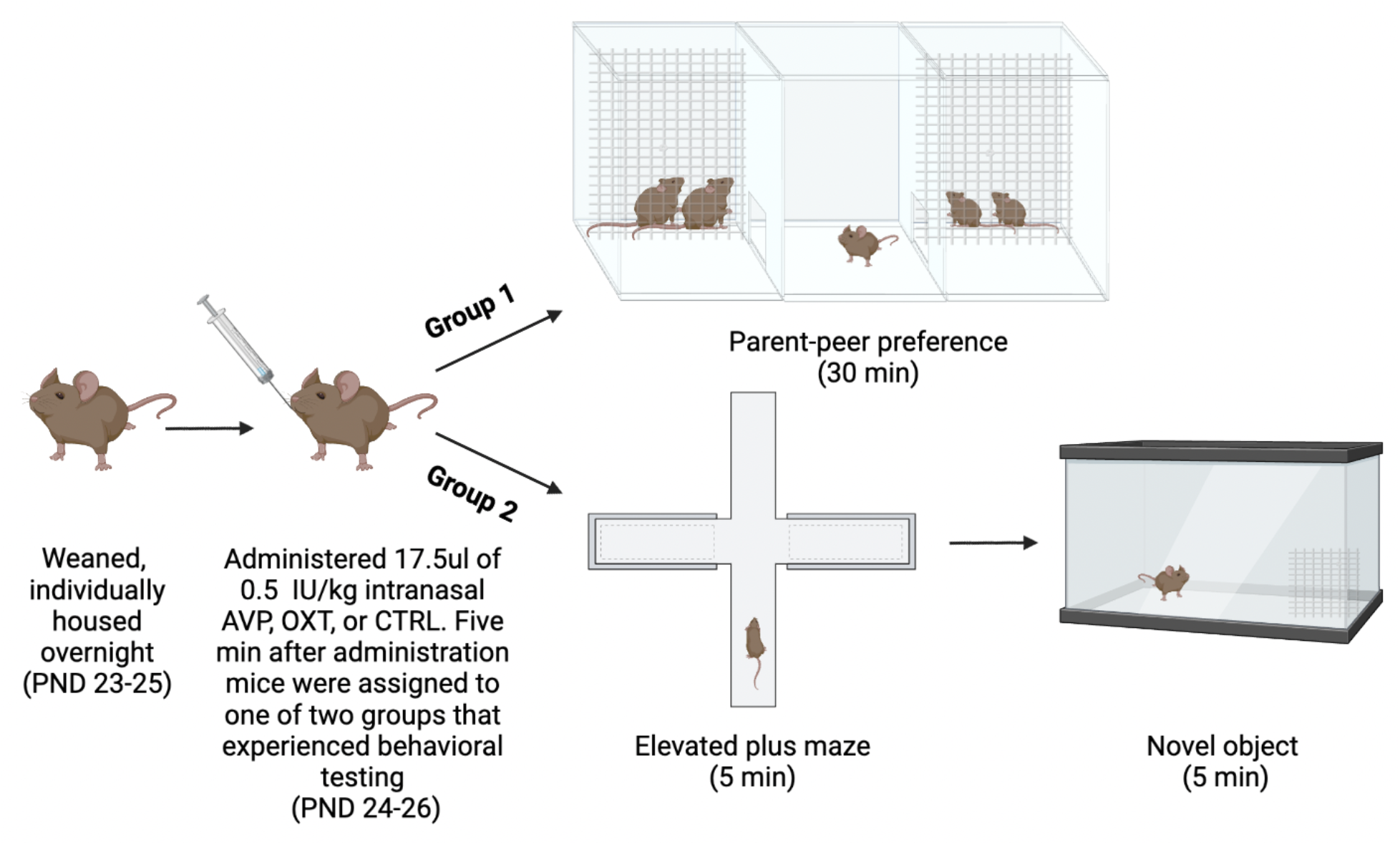
Timeline and schematic for experimental design. All mice were weaned from their home cage, housed individually overnight, and randomly assigned an intranasal treatment. Approximately half of the mice were assigned to Group 1, and the other half were assigned to Group 2.

#### Parent-peer preference test

Each Group 1 mouse was tested in the parent-peer preference test using a three-chambered apparatus (91 cm x 46 cm x 43 cm) divided into three equal chambers. Each side chamber had a wire mesh partition at the back (30 cm x 10 cm) where stimulus animals could be presented. Stimulus mice were as follows: the mother and father mice were placed on one side of the three-chambered Plexiglas cage behind the wire mesh, and a pair of age-matched female and male sibling mice (PND 24-31) to the focal mouse and its parents were placed on the other side of the apparatus. Similar tests have been previously used to assess social preferences in California mice in the Marler laboratory (Zhao & Marler, 2014; Zhao & Marler, 2016; Zhao et al., 2020) and in other labs to assess social preferences such as partner preference (Williams et al., 1992; Bales & Carter, 2003) and peer preference (Beery et al., 2018; Lee & Beery, 2021). Before each test, the parents of the focal mouse were randomly assigned to either the left or right side of the chamber, and unrelated, age-matched peers (one female, one male) were placed on the other side of the chamber. Once stimulus animals were placed in their respective chambers in the testing apparatus, each peri-adolescent mouse was given their randomly assigned IN dose (AVP, OXT, or CTRL) in their singly housed home cage. Five minutes after IN administration, the test mouse was placed in the center chamber of the testing apparatus, and its behavior was videotaped for 30 min. After this test, peri-adolescent mice were euthanized, and their brains were extracted for future studies.

#### Elevated plus maze test and novel object test

Mice assigned to Group 2 were first given the elevated plus maze test, followed by a novel object test. The maze consisted of two open and two enclosed opaque arms, each 67 cm long and 5.5 cm wide. The arms were elevated 1 m above the floor. Peri-adolescent test mice were given IN treatment in their home cage five minutes prior to behavioral testing. At the start of the test, each mouse was placed in the center of the maze, and its behavior was videotaped for five min. Any animals that jumped off the open arms of the maze were captured and placed back in the center of the maze. Throughout the study, six mice fell off the apparatus, one time each. Immediately following the elevated plus maze test, mice were moved to a new testing room and placed on the far side of a glass arena (50cm x 30cm x 30cm) that contained a novel 5cm x 5cm x 5cm metal cube, and behavior was recorded for five min. Peri-adolescent mice were then euthanized, and their brains were extracted for future studies.

### Behavior Quantification

For an ethogram describing each behavior measured, see **S. Table 1**. For each behavior, we used a continuous sampling method. All experimenters were blind to treatment conditions during quantification.

#### Parent-peer preference test

For the trial to be successful, peri-adolescent mice had to visit both sides of the chamber in the first 10 min of the test to be scored. Two male mice did not meet this threshold and were excluded because they did not enter the chamber with the two peer mice. Time to enter each chamber and time spent in each of the three chambers were scored.

#### Elevated plus maze test

Trained observers scored behavior live during the test and recorded the duration of time in the open arms and the number of crosses through the center of the maze.

#### Novel object test

Behavior videos were scored for latency to approach the novel object, and time spent investigating the novel object was measured by the total number of seconds.

### Data Analysis

ANOVA tests were conducted for each behavioral test to compare saline control, OXT, and AVP treatment outcomes. The significance level was set at p < 0.05 for all analyses, and all tests were two-tailed. All reported p-values were corrected using Benjamini-Hochberg false discovery rate corrections to control for multiple comparisons when the effect of an X variable was tested for a relationship with multiple Y variables.

## RESULTS

### Parent-peer preference test

We conducted a three-chambered social preference test to assess the effects of an acute pulse of OXT or AVP on juvenile social preference for their parents versus novel peers. Regardless of treatment, juvenile females preferred their parents over their peers (*F*_1,22_=25.09, *p<*0.0001], and their peers over the empty middle chamber (*F*_1,22_=55.38, *p<*0.0001) (**Fig. 2A**). Likewise, regardless of treatment, juvenile males preferred their parents over their peers (*F*_1,20_=37.16, *p*<0.0001), and their peers over the empty chamber (*F*_1,20_=25.15, *p*<0.0001) (**Fig. 2B**). To test for the effects of IN OXT and AVP on social preference, we created a preference score by subtracting the total percentage of each individual’s preference for their peers from their preference for their parents [(time spent with parents/total test time)-(time spent with peers/total test time)]. There were no effects of treatment on the social preference expressed by females (*F*_2, 20_=1.38, *p*=0.27) (**Fig. 2C**). However, IN OXT increased juvenile male preference for their parents (*F*_2, 18_=4.98, *p*<0.05, ΔR^2^=0.3562) (**Fig. 2D**). There were no effects of treatment on preference for peers over an empty chamber expressed in females (*F*_2, 20_=0.56, *p*=0.58) or males (*F*_2, 18_=0.16, *p*=0.85) (**Fig. 2E-F**).

**Figure 2.**
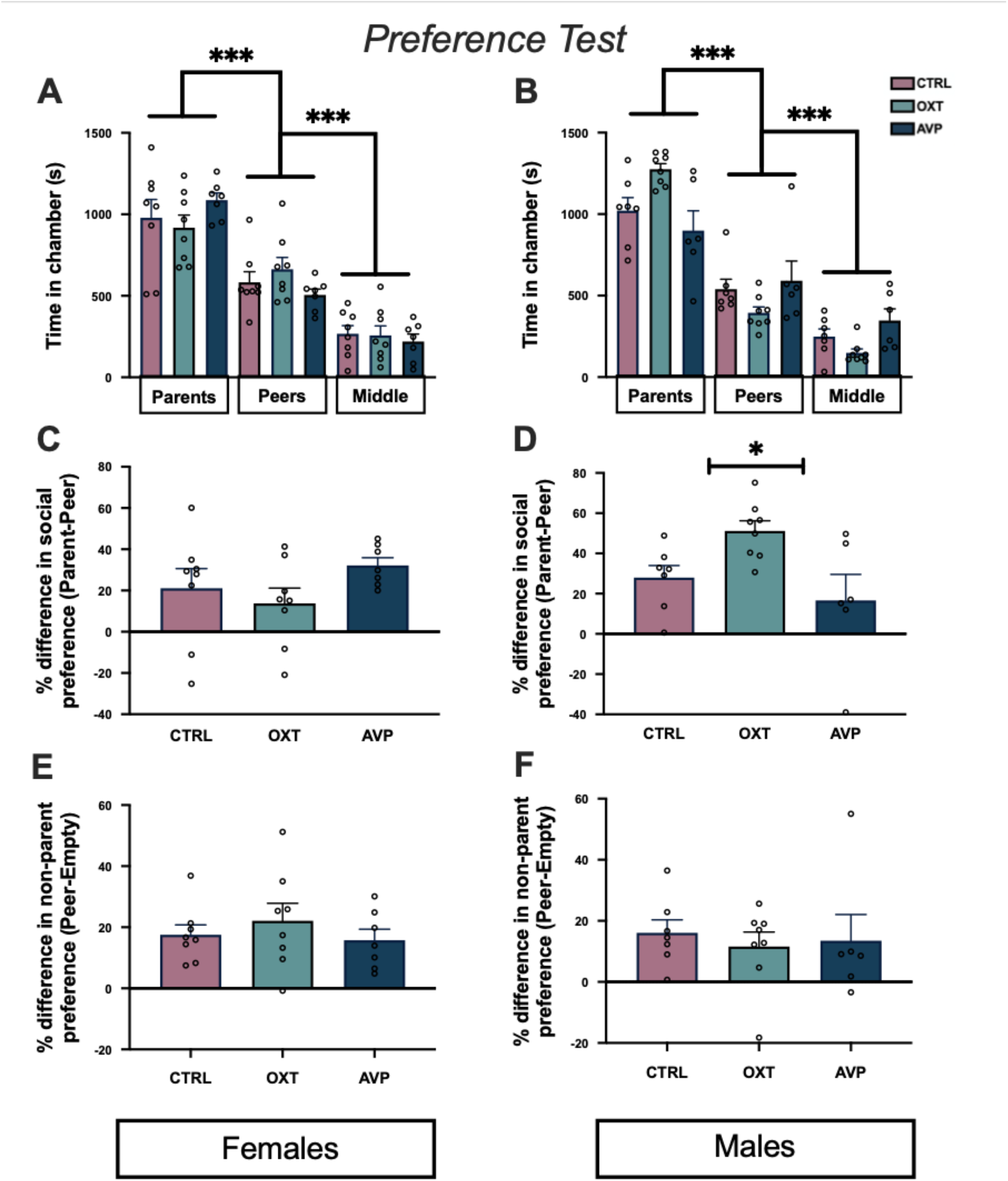
(**A**) Both female and (**B**) male juveniles preferred their parents compared to their peers or the empty chamber and preferred their peers over an empty chamber. (**C**) Neither IN OXT nor AVP influenced social preference in female juveniles. (**D**) IN OXT increased juvenile male preference for their parents above and beyond their natural preference for their parents. (**E**) Neither IN OXT nor AVP influenced preference for peers over an empty chamber in female juveniles. (**F**) Neither IN OXT nor AVP influenced preference for peers over an empty chamber in male juveniles.

### Elevated plus maze test

The goal was to assess the effects of OXT or AVP on juvenile exploration. There were no treatment effects of time spent on the open arms for females (*F*_2,26_=0.31, *p*=0.74) or males (*F*_2,23_=1.37, *p*=0.27) (**Fig. 3A**). There were also no treatment effects on the number of crosses through the center of the apparatus for females (*F*_2,26_=0.20, *p*=0.82) or males (*F*_2,23_=0.47, *p*=0.63) (**Fig. 3B**).

**Figure 3.**
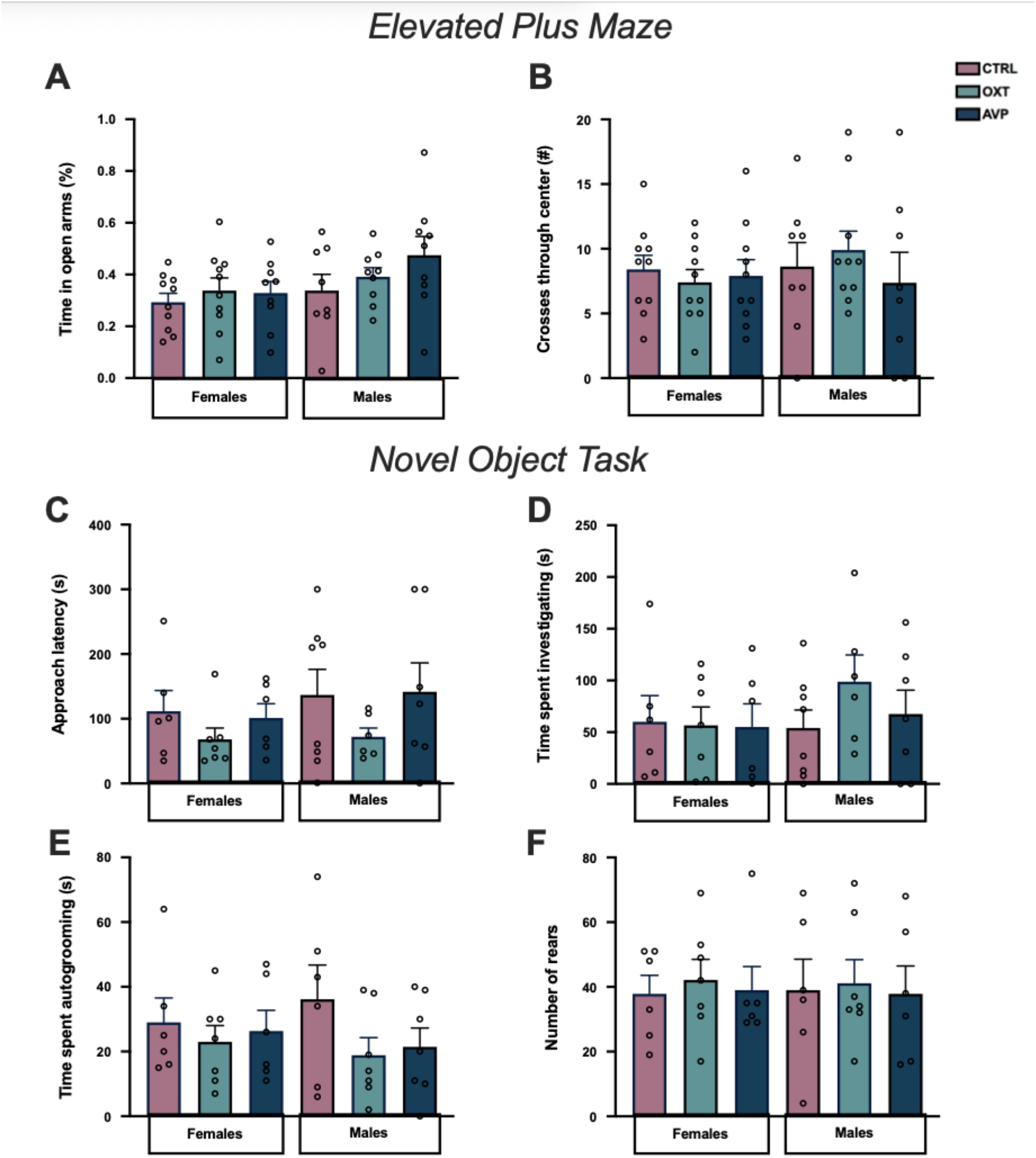
*Elevated Plus Maze*. Neither IN OXT nor AVP influenced (**A**) time spent on open arms (**B**) or the number of crosses through the center of the elevated plus maze. *Novel Object*. Neither IN OXT nor IN AVP influenced (**C**) latency to approach the novel object, (**D**) time spent investigating the novel object, (**E**) self-grooming, or (**F**) locomotor activity.

### Novel object test

We measured behavior in a novel object task to assess the effects of an acute pulse of OXT or AVP on juvenile response to novelty. There were no treatment effects of latency to approach the novel object for females (*F*_2,16_=0.94, *p*=0.41) or males (*F*_2,18_=0.98, *p*=0.40) (**Fig. 3C**). There were also no treatment effects of time spent interacting with the novel object for females (*F*_2,16_=0.01, *p*=0.99) or males (*F*_2,18_=1.06, *p*=0.37) (**Fig. 3D**). There were also no differences in self-grooming for females (*F*_2,16_=0.23, *p*=0.79) or males (*F*_2,18_=1.56, *p*=0.24) (**Fig. 3E**). Finally, there were no differences in locomotor activity for females (*F*_2,16_=0.12, *p*=0.89) or males (*F*_2,18_=0.04, *p*=0.96) (**Fig. 3F**).

## DISCUSSION

Adolescence is a time of profound physiological and social change; however, the neural underpinnings of these social changes are just beginning to be explored (Pfeifer & Allen, 2021). To our knowledge, this is the first study to test the influence of a hormonal substrate on social preferences for parents versus peers in peri-adolescents. This study found that OXT but not AVP increased social preference in peri-adolescent males for their parents. Female peri-adolescents did not change their social preferences in response to either OXT or AVP, suggesting that other mechanisms likely have greater impacts on female social preferences during peri-adolescence. Because neither OXT nor AVP influenced performance in the elevated plus maze task nor the novel object task, we argue that OXT has an effect specific to social behavior rather than sensation-seeking, which is also heightened during the peri-adolescent window in many species (Arnet, 1996; Stansfield et al., 2004).

The results raise two issues which are: why is there a sex difference, and why is only OXT influencing behavior? While female and male California mice are very similar in their behaviors, one notable ecological difference occurs in behavior around the time of adolescence; females disperse farther from the natal territory than males (Ribble, 1992). Males can even take over their father’s territory (personal communication, R. Petric). Anecdotal evidence in our lab suggests that when weaning a cage of juveniles at postnatal day 28, males are more likely to be still suckling on their mothers than females, possibly providing future insight into the sex differences we observed in this study. We speculate that males may require more plasticity in mechanisms influencing the parent-offspring bond, such as continued responsiveness to oxytocin to allow adjustments in social tolerance or bonding. However, species differences should also be considered when interpreting the sex differences found in this study. In contrast to California mice, the males of most other mammalian species are the ones that typically disperse farther from the natal territory (Pocock et al., 2005; Thompson, 2009). Therefore, it is possible that this effect of OXT is specific to California mice or mammals with a similar dispersal pattern.

Regardless of the reason for why there were sex differences in the response of male and female juveniles to OXT, there are several speculative reasons why OXT made peri-adolescent males prefer to spend more time with their parents. One possibility is that IN OXT administration reinforces the parent-offspring social bond. In this study, the peri-adolescents were removed from their home cage for 24 hours prior to testing; the loss of physical contact with their natal nest and parents could have caused OXT levels to decrease by removing the touch, smell, and gaze components of the social bond that reinforce the positive feedback release of OXT (Neumann et al., 1996; Romero et al., 2014; Wang et al., 2022), and IN OXT could have reinstated this positive feedback signaling. Studies in prairie voles have shown that adult reproductive and non-reproductive social behavior can be altered by the quality of the natal rearing, suggesting that the parent-offspring social interactions meaningfully impact the brain and behavior (Ahern et al., 2021; Perkeybile et al., 2019; Perkeybile & Bales, 2017; Bales et al., 2018). More broadly, for males, OXT and familiarity may be driving the heightened preference. OXT can increase the memory of and attention and affection towards familiar individuals (Marsh et al., 2021; Rimmele et al., 2009; Lu et al., 2019; Sheele et al., 2013). Therefore, we cannot rule out the possibility that familiarity is driving social preference for their parents. However, if the change in social preference is driven by familiarity, it is unlikely to be caused by anxiety because OXT did not influence elevated plus behavior. This is consistent with a study in prairie voles that found philopatry was not linked to high anxiety (del Razo & Bales, 2016). It is also unlikely that novelty avoidance led to the change in social preference because there was no difference in novel object task performance and the OXT male mice spent ∼43% more time in the peer chamber than in the empty chamber where they could avoid novel social contact. This increase is similar to control-treated males (∼46%). For the above reasons, we argue that OXT is enhancing prosocial behavior on the basis of the offspring-parent bond and/or familiarity.

Again, we can only speculate about the sex differences and why AVP is not influencing behavior during peri-adolescence. It is, however, interesting to note that adult male, but not female, California mice have greater OXT receptor binding in the cingulate cortex and bed nucleus of the stria terminalis, two areas known for emotion processing and social behavior. In contrast, there is no difference in AVP receptor binding (Insel et al., 1991). Sex differences in response to neuropeptides are often found, but it has been difficult to identify why this occurs. As the functions of neuropeptides during peri-adolescence are understudied, more research will be needed to examine functions, especially since neuropeptide effects on behavior can be context-dependent (see Reviews by Carter et al., 2020; Caldwell, 2018; Bartz et al., 2011; Rieger et al., 2022).

The neurobiological underpinnings of social bond maintenance in peri-adolescence are important to study because they offer a unique window into the factors associated with changing social preferences. Mammalian offspring are born and immediately initiate a social bond with their mother and possibly both parents in bi-parental species. However, the maintenance of these social bonds is tentative and depends on experience, environmental state, and biological substrate signaling. Most mammals, including humans, will start to lose their social preference for their parents over time. In peri-adolescent humans, activation of the nucleus accumbens to the ventromedial prefrontal cortex circuit is dependent on age: in younger adolescents (∼7-12 years old), it is activated in response to their mother’s voice, but in older adolescents (∼13-16 years old) it is activated by the voices of strangers (Abrams et al., 2022). Because the nucleus accumbens to the ventromedial prefrontal cortex is known for both reward and social evaluation, these results underscore the importance of social sensory signals triggering the expression of social preferences.

This study is the first to demonstrate that OXT influences the maintenance of offspring-parent social preferences during peri-adolescence in males but not females. As such, this finding suggests that sex differences may also play a role in the mechanisms underlying social bond maintenance within family units. This work adds to the growing body of literature on social bonds within family units and with unrelated conspecifics.

## ACKNOWLEDGMENTS

We would like to thank NSF grant IOS-1946613 for generously funding this research, Anna Backenger, Ekenedilichukwu Ikegwuani, and Lauryn Novak for their help with behavioral experiments and the UW Madison Animal Care Staff for their excellent care of the mice, and, finally, the Serendipity Scholarship Award for summer funding.

## REFERENCES

Abrams, D. A., Mistry, P. K., Baker, A. E., Padmanabhan, A., & Menon, V. (2022). A neurodevelopmental shift in reward circuitry from mother’s to nonfamilial voices in adolescence. Journal of Neuroscience, 42(20), 4164–4173.

Acevedo-Rodriguez, A., Mani, S. K., & Handa, R. J. (2015). Oxytocin and estrogen receptor β in the brain: an overview. Frontiers in Endocrinology, 6, 160.

Ahern, T. H., Olsen, S., Tudino, R., & Beery, A. K. (2021). Natural variation in the oxytocin receptor gene and rearing interact to influence reproductive and nonreproductive social behavior and receptor binding. Psychoneuroendocrinology, 128, 105209.

Albers, H. E. (2012). The regulation of social recognition, social communication and aggression: vasopressin in the social behavior neural network. Hormones and Behavior, 61(3), 283–292.

Albers, H. E., & Bamshad, M. (1999). Role of vasopressin and oxytocin in the control of social behavior in Syrian hamsters (Mesocricetus auratus. Progress in Brain Research, 119, 395–408.

Anagnostou, E., Soorya, L., Brian, J., Dupuis, A., Mankad, D., Smile, S., & Jacob, S. (2014). Intranasal oxytocin in the treatment of autism spectrum disorders: a review of literature and early safety and efficacy data in youth. Brain Research, 1580, 188–198.

Arnett, J. J. (1996). Sensation seeking, aggressiveness, and adolescent reckless behavior. Personality and Individual Differences, 20(6), 693–702.

Bales, K. L., & Carter, C. S. (2003). Sex differences and developmental effects of oxytocin on aggression and social behavior in prairie voles (Microtus ochrogaster). Hormones and Behavior, 44(3), 178–184.

Bales, K.L., Witczak, L.R., Simmons, T.C., Savidge, L.E., Rothwell, E.S., Rogers, F.D., Manning, R.A., Heise, M.J., Englund, M. and Del Razo, R.A. (2018). Social touch during development: Long-term effects on brain and behavior. Neuroscience & Biobehavioral Reviews, 95, 202–219.

Bartz, J. A., Zaki, J., Bolger, N., & Ochsner, K. N. (2011). Social effects of oxytocin in humans: context and person matter. Trends in Cognitive Sciences, 15(7), 301–309.

Baxter, A., Anderson, M., Seelke, A. M., Kinnally, E. L., Freeman, S. M., & Bales, K. L. (2020). Oxytocin receptor binding in the titi monkey hippocampal formation is associated with parental status and partner affiliation. Scientific Reports, 10(1), 1–14.

Bechard, A., & Mason, G. (2010). Leaving home: a study of laboratory mouse pup independence. Applied Animal Behaviour Science, 125(3-4), 181–188.

Beery, A. K., Christensen, J. D., Lee, N. S., & Blandino, K. L. (2018). Specificity in sociality: mice and prairie voles exhibit different patterns of peer affiliation. Frontiers in Behavioral Neuroscience, 12, 50.

Bester-Meredith, J. K., & Marler, C. A. (2001). Vasopressin and aggression in cross-fostered California mice (Peromyscus californicus) and white-footed mice (Peromyscus leucopus). Hormones and Behavior, 40(1), 51–64.

Bredewold, R., & Veenema, A. H. (2018). Sex differences in the regulation of social and anxiety-related behaviors: insights from vasopressin and oxytocin brain systems. Current Opinion in Neurobiology, 49, 132–140.

Brenhouse, H. C., & Andersen, S. L. (2011). Developmental trajectories during adolescence in males and females: a cross-species understanding of underlying brain changes. Neuroscience & Biobehavioral Reviews, 35(8), 1687–1703.

Caldwell, H. K. (2018). Oxytocin and sex differences in behavior. Current opinion in behavioral sciences, 23, 13–20.

Carter, C.S., Kenkel, W.M., MacLean, E.L., Wilson, S.R., Perkeybile, A.M., Yee, J.R., Ferris, C.F., Nazarloo, H.P., Porges, S.W., Davis, J.M. and Connelly, J.J. (2020). Is oxytocin “nature’s medicine”. Pharmacological Reviews, 72(4), 829–861.

Carter, C. S., Williams, J. R., Witt, D. M., & Insel, T. R. (1992). Oxytocin and social bonding. Annals of the New York Academy of Sciences, 652, 204–211.

Crockford, C., Wittig, R. M., Langergraber, K., Ziegler, T. E., Zuberbühler, K., & Deschner, T. (2013). Urinary oxytocin and social bonding in related and unrelated wild chimpanzees. Proceedings of the Royal Society B: Biological Sciences, 280(1755), 20122765.

Curley, J. P., Jensen, C. L., Franks, B., & Champagne, F. A. (2012). Variation in maternal and anxiety-like behavior associated with discrete patterns of oxytocin and vasopressin 1a receptor density in the lateral septum. Hormones and Behavior, 61(3), 454–461.

de Moura Oliveira, V. E., Neumann, I. D., & de Jong, T. R. (2019). Post-weaning social isolation exacerbates aggression in both sexes and affects the vasopressin and oxytocin system in a sexspecific manner. Neuropharmacology, 156, 107504.

Delevich, K., Klinger, M., Okada, N. J., & Wilbrecht, L. (2021). Coming of age in the frontal cortex: the role of puberty in cortical maturation. In Seminars in Cell & Developmental Biology (Vol. 118, pp. 64–72). Academic Press.

Del Razo, R. A., & Bales, K. L. (2016). Exploration in a dispersal task: effects of early experience and correlation with other behaviors in prairie voles (Microtus ochrogaster). Behavioural Processes, 132, 66–75.

De Vries, G. J., Wang, Z., Bullock, N. A., & Numan, S. (1994). Sex differences in the effects of testosterone and its metabolites on vasopressin messenger RNA levels in the bed nucleus of the stria terminalis of rats. Journal of Neuroscience, 14(3), 1789–1794.

Duell, N., & Steinberg, L. (2019). Positive risk-taking in adolescence. Child Development Perspectives, 13(1), 48–52.

Duque-Wilckens, N., Steinman, M.Q., Busnelli, M., Chini, B., Yokoyama, S., Pham, M., Laredo, S.A., Hao, R., Perkeybile, A.M., Minie, V.A. and Tan, P.B. (2018). Oxytocin receptors in the anteromedial bed nucleus of the stria terminalis promote stress-induced social avoidance in female California mice. Biological Psychiatry, 83(3), 203–213.

Febo, M., Numan, M., & Ferris, C. F. (2005). Functional magnetic resonance imaging shows oxytocin activates brain regions associated with mother–pup bonding during suckling. Journal of Neuroscience, 25(50), 11637–11644.

Forbes, E. E., & Dahl, R. E. (2010). Pubertal development and behavior: hormonal activation of social and motivational tendencies. Brain and Cognition, 72(1), 66–72.

Frazier, C. R., Trainor, B. C., Cravens, C. J., Whitney, T. K., & Marler, C. A. (2006). Paternal behavior influences development of aggression and vasopressin expression in male California mouse offspring. Hormones and Behavior, 50(5), 699–707.

Frayne, J., & Nicholson, H. D. (1995). Effect of oxytocin on testosterone production by isolated rat Leydig cells is mediated via a specific oxytocin receptor. Biology of Reproduction, 52(6), 1268–1273.

Freeman, S. M., Samineni, S., Allen, P. C., Stockinger, D., Bales, K. L., Hwa, G. G., & Roberts, J. A. (2016). Plasma and CSF oxytocin levels after intranasal and intravenous oxytocin in awake macaques. Psychoneuroendocrinology, 66, 185–194.

Gubernick, D. J., Winslow, J. T., Jensen, P., Jeanotte, L., & Bowen, J. (1995). Oxytocin changes in males over the reproductive cycle in the monogamous, biparental California mouse, Peromyscus californicus. Hormones and Behavior, 29(1), 59–73.

Guoynes, C. D., & Marler, C. A. (2021). An acute dose of intranasal oxytocin rapidly increases maternal communication and maintains maternal care in primiparous postpartum California mice. Plos One, 16(4), e0244033.

Guoynes, C. D., & Marler, C. A. (2022). Intranasal oxytocin reduces pre-courtship aggression and increases paternal response in California mice (Peromyscus californicus). Physiology & Behavior, 249, 113773.

Guoynes, C. D., Simmons, T. C., Downing, G. M., Jacob, S., Solomon, M., & Bales, K. L. (2018). Chronic intranasal oxytocin has dose-dependent effects on central oxytocin and vasopressin systems in prairie voles (Microtus ochrogaster). Neuroscience, 369, 292–302.

Heinrichs, M., von Dawans, B., & Domes, G. (2009). Oxytocin, vasopressin, and human social behavior. Frontiers in Neuroendocrinology, 30(4), 548–557.

Horii-Hayashi, N., Sasagawa, T., Matsunaga, W., Matsusue, Y., Azuma, C., & Nishi, M. (2013). Developmental changes in desensitisation of c-Fos expression induced by repeated maternal separation in pre-weaned mice. Journal of Neuroendocrinology, 25(2), 158–167.

Huang, Y., Huang, X., Ebstein, R. P., & Yu, R. (2021). Intranasal oxytocin in the treatment of autism spectrum disorders: A multilevel meta-analysis. Neuroscience & Biobehavioral Reviews, 122, 18–27.

Insel, T. R. (1990). Regional changes in brain oxytocin receptors post-partum: time-course and relationship to maternal behaviour. Journal of Neuroendocrinology, 2(4), 539–545.

Insel, T. R., Gelhard, R., & Shapiro, L. E. (1991). The comparative distribution of forebrain receptors for neurohypophyseal peptides in monogamous and polygamous mice. Neuroscience, 43(2-3), 623–630.

Jarcho, M. R., Mendoza, S. P., Mason, W. A., Yang, X., & Bales, K. L. (2011). Intranasal vasopressin affects pair bonding and peripheral gene expression in male Callicebus cupreus. Genes, Brain and Behavior, 10(3), 375–383.

Keebaugh, A. C., Barrett, C. E., LaPrairie, J. L., Jenkins, J. J., & Young, L. J. (2015). RNAi knockdown of oxytocin receptor in the nucleus accumbens inhibits social attachment and parental care in monogamous female prairie voles. Social Neuroscience, 10(5), 561–570.

Kendrick, K. M. (2000). Oxytocin, motherhood and bonding. Experimental Physiology, 85(1), 111s–124s.

Kent, K., Arientyl, V., Khachatryan, M. M., & Wood, R. I. (2013). Oxytocin induces a conditioned social preference in female mice. Journal of Neuroendocrinology, 25(9), 803–810.

Keverne, E. B., & Curley, J. P. (2004). Vasopressin, oxytocin and social behaviour. Current Opinion in Neurobiology, 14(6), 777–783.

Keverne, E. B., & Kendrick, K. M. (1992). Oxytocin Facilitation of Maternal Behavior in Sheep. Annals of the New York Academy of Sciences, 652(1), 83–101.

Koolschijn, P. C. M., Peper, J. S., & Crone, E. A. (2014). The influence of sex steroids on structural brain maturation in adolescence. PloS One, 9(1), e83929.

Lee, M.R., Scheidweiler, K.B., Diao, X.X., Akhlaghi, F., Cummins, A., Huestis, M.A., Leggio, L. and Averbeck, B.B. (2018). Oxytocin by intranasal and intravenous routes reaches the cerebrospinal fluid in rhesus macaques: determination using a novel oxytocin assay. Molecular Psychiatry, 23(1), 115–122.

Lee, M.R., Shnitko, T.A., Blue, S.W., Kaucher, A.V., Winchell, A.J., Erikson, D.W., Grant, K.A. and Leggio, L. (2020). Labeled oxytocin administered via the intranasal route reaches the brain in rhesus macaques. Nature Communications, 11(1), 1–10.

Lee, N. S., & Beery, A. K. (2021). The role of dopamine signaling in prairie vole peer relationships. Hormones and Behavior, 127, 104876.

Liu, Y., & Wang, Z. X. (2003). Nucleus accumbens oxytocin and dopamine interact to regulate pair bond formation in female prairie voles. Neuroscience, 121(3), 537–544.

Loth, M. K., & Donaldson, Z. R. (2021). Oxytocin, dopamine, and opioid interactions underlying pair bonding: highlighting a potential role for microglia. Endocrinology, 162(2), bqaa223.

Lu, Q., Lai, J., Du, Y., Huang, T., Prukpitikul, P., Xu, Y., & Hu, S. (2019). Sexual dimorphism of oxytocin and vasopressin in social cognition and behavior. Psychology Research and Behavior Management, 12, 337.

Lukas, M., & Neumann, I. D. (2014). Social preference and maternal defeat-induced social avoidance in virgin female rats: sex differences in involvement of brain oxytocin and vasopressin. Journal of Neuroscience Methods, 234, 101–107.

Lukas, M., Toth, I., Reber, S. O., Slattery, D. A., Veenema, A. H., & Neumann, I. D. (2011). The neuropeptide oxytocin facilitates pro-social behavior and prevents social avoidance in rats and mice. Neuropsychopharmacology, 36(11), 2159–2168.

Marsh, N., Scheele, D., Postin, D., Onken, M., & Hurlemann, R. (2021). Eye-Tracking reveals a role of oxytocin in attention allocation towards familiar faces. Frontiers in Endocrinology, 12, 629760.

Maud, C., Ryan, J., McIntosh, J. E., & Olsson, C. A. (2018). The role of oxytocin receptor gene (OXTR) DNA methylation (DNAm) in human social and emotional functioning: a systematic narrative review. BMC Psychiatry, 18(1), 1–13.

McCarthy, M. M., McDonald, C. H., Brooks, P. J., & Goldman, D. (1996). An anxiolytic action of oxytocin is enhanced by estrogen in the mouse. Physiology & Behavior, 60(5), 1209–1215.

Monari, P. K., Rieger, N. S., Schefelker, J., & Marler, C. A. (2021). Intranasal oxytocin drives coordinated social approach. Scientific Reports, 11(1), 1–13.

Murakami, G., Hunter, R. G., Fontaine, C., Ribeiro, A., & Pfaff, D. (2011). Relationships among estrogen receptor, oxytocin and vasopressin gene expression and social interaction in male mice. European Journal of Neuroscience, 34(3), 469–477.

Nagasawa, M., Okabe, S., Mogi, K., & Kikusui, T. (2012). Oxytocin and mutual communication in mother-infant bonding. Frontiers in Human Neuroscience, 6, 31.

Neumann, I., Douglas, A. J., Pittman, Q. J., Russell, J. A., & Landgraf, R. (1996). Oxytocin released within the supraoptic nucleus of the rat brain by positive feedback action is involved in parturition-related events. Journal of Neuroendocrinology, 8(3), 227–233.

Neumann, I., Ludwig, M., Engelmann, M., Pittman, Q. J., & Landgraf, R. (1993). Simultaneous microdialysis in blood and brain: oxytocin and vasopressin release in response to central and peripheral osmotic stimulation and suckling in the rat. Neuroendocrinology, 58(6), 637–645.

Neumann, I. D., Maloumby, R., Beiderbeck, D. I., Lukas, M., & Landgraf, R. (2013). Increased brain and plasma oxytocin after nasal and peripheral administration in rats and mice. Psychoneuroendocrinology, 38(10), 1985–1993.

Nishimori, K., Young, L. J., Guo, Q., Wang, Z., Insel, T. R., & Matzuk, M. M. (1996). Oxytocin is required for nursing but is not essential for parturition or reproductive behavior. Proceedings of the National Academy of Sciences, 93(21), 11699–11704.

Oppong-Damoah, A., Zaman, R. U., D’Souza, M. J., & Murnane, K. S. (2019). Nanoparticle encapsulation increases the brain penetrance and duration of action of intranasal oxytocin. Hormones and Behavior, 108, 20–29.

Pedersen, C. A., Caldwell, J. D., Walker, C., Ayers, G., & Mason, G. A. (1994). Oxytocin activates the postpartum onset of rat maternal behavior in the ventral tegmental and medial preoptic areas. Behavioral Neuroscience, 108(6), 1163.

Peper, J. S., Pol, H. H., Crone, E. A., & Van Honk, J. (2011). Sex steroids and brain structure in pubertal boys and girls: a mini-review of neuroimaging studies. Neuroscience, 191, 28–37.

Perea-Rodriguez, J. P., Takahashi, E. Y., Amador, T. M., Hao, R. C., Saltzman, W., & Trainor, B. C. (2015). Effects of reproductive experience on central expression of progesterone, oestrogen α, oxytocin and vasopressin receptor mRNA in male California mice (Peromyscus californicus). Journal of Neuroendocrinology, 27(4), 245–252.

Perkeybile, A. M., & Bales, K. L. (2017). Intergenerational transmission of sociality: the role of parents in shaping social behavior in monogamous and non-monogamous species. Journal of Experimental Biology, 220(1), 114–123.

Perkeybile, A.M., Carter, C.S., Wroblewski, K.L., Puglia, M.H., Kenkel, W.M., Lillard, T.S., Karaoli, T., Gregory, S.G., Mohammadi, N., Epstein, L. and Bales, K.L., 2019. Early nurture epigenetically tunes the oxytocin receptor. Psychoneuroendocrinology, 99, pp.128–136.

Pfeifer, J. H., & Allen, N. B. (2021). Puberty initiates cascading relationships between neurodevelopmental, social, and internalizing processes across adolescence. Biological Psychiatry, 89(2), 99–108.

Pocock, M. J., Hauffe, H. C., & Searle, J. B. (2005). Dispersal in house mice. Biological Journal of the Linnean Society, 84(3), 565–583.

Prounis, G. S., Thomas, K., & Ophir, A. G. (2018). Developmental trajectories and influences of environmental complexity on oxytocin receptor and vasopressin 1A receptor expression in male and female prairie voles. Journal of Comparative Neurology, 526(11), 1820–1842.

Prugh, L. R., Deguines, N., Grinath, J. B., Suding, K. N., Bean, W. T., Stafford, R., & Brashares, J. S. (2018). Ecological winners and losers of extreme drought in California. Nature Climate Change, 8(9), 819–824.

Ribble, D. O. (1992). Dispersal in a monogamous rodent, Peromyscus californicus. Ecology, 73(3), 859–866.

Rieger, N. S., Guoynes, C. D., Monari, P. K., Hammond, E. R., Malone, C. L., & Marler, C. A. (2022). Neuroendocrine mechanisms of aggression in rodents. Motivation Science, 8(2).

Rimmele, U., Hediger, K., Heinrichs, M., & Klaver, P. (2009). Oxytocin makes a face in memory familiar. Journal of Neuroscience, 29(1), 38–42.

Romeo, R. D. (2003). Puberty: a period of both organizational and activational effects of steroid hormones on neurobehavioural development. Journal of neuroendocrinology, 15(12), 1185–1192.

Romero, T., Nagasawa, M., Mogi, K., Hasegawa, T., & Kikusui, T. (2014). Oxytocin promotes social bonding in dogs. Proceedings of the National Academy of Sciences, 111(25), 9085–9090.

Scheele, D., Wille, A., Kendrick, K.M., Stoffel-Wagner, B., Becker, B., Güntürkün, O., Maier, W. and Hurlemann, R. (2013). Oxytocin enhances brain reward system responses in men viewing the face of their female partner. Proceedings of the National Academy of Sciences, 110(50), 20308–20313.

Shapiro, R. A., Xu, C., & Dorsa, D. M. (2000). Differential transcriptional regulation of rat vasopressin gene expression by estrogen receptor α and β. Endocrinology, 141(11), 4056–4064.

Skuse, D.H., Lori, A., Cubells, J.F., Lee, I., Conneely, K.N., Puura, K., Lehtimäki, T., Binder, E.B. and Young, L.J. (2014). Common polymorphism in the oxytocin receptor gene (OXTR) is associated with human social recognition skills. Proceedings of the National Academy of Sciences, 111(5), 1987–1992.

Smith, A. S., Korgan, A. C., & Young, W. S. (2019). Oxytocin delivered nasally or intraperitoneally reaches the brain and plasma of normal and oxytocin knockout mice. Pharmacological Research, 146, 104324.

Smith, C. J., Poehlmann, M. L., Li, S., Ratnaseelan, A. M., Bredewold, R., & Veenema, A. H. (2017). Age and sex differences in oxytocin and vasopressin V1a receptor binding densities in the rat brain: focus on the social decision-making network. Brain Structure and Function, 222(2), 981–1006.

Spear, L. (2007). The Developing Brain and Adolescent-Typical Behavior Patterns An Evolutionary Approach. Adolescent Psychopathology and the Developing Brain: Integrating Brain and Prevention Science, 9.

Staes, N., Koski, S. E., Helsen, P., Fransen, E., Eens, M., & Stevens, J. M. (2015). Chimpanzee sociability is associated with vasopressin (Avpr1a) but not oxytocin receptor gene (OXTR) variation. Hormones and Behavior, 75, 84–90.

Stansfield, K. H., Philpot, R. M., & Kirstein, C. L. (2004). An animal model of sensation seeking: the adolescent rat. Annals of the New York Academy of Sciences, 1021(1), 453–458.

Steinberg, L. (2010). A dual systems model of adolescent risk-taking. Developmental Psychobiology: The Journal of the International Society for Developmental Psychobiology, 52(3), 216–224.

Steinman, M. Q., Duque-Wilckens, N., & Trainor, B. C. (2019). Complementary neural circuits for divergent effects of oxytocin: social approach versus social anxiety. Biological Psychiatry, 85(10), 792–801.

Steinman, M.Q., Duque-Wilckens, N., Greenberg, G.D., Hao, R., Campi, K.L., Laredo, S.A., Laman-Maharg, A., Manning, C.E., Doig, I.E., Lopez, E.M. and Walch, K. (2016). Sex-specific effects of stress on oxytocin neurons correspond with responses to intranasal oxytocin. Biological Psychiatry, 80(5), 406–414.

Striepens, N., Kendrick, K. M., Hanking, V., Landgraf, R., Wüllner, U., Maier, W., & Hurlemann, R. (2013). Elevated cerebrospinal fluid and blood concentrations of oxytocin following its intranasal administration in humans. Scientific Reports, 3(1), 1–5.

Taylor, J. H., Cavanaugh, J., & French, J. A. (2017). Neonatal oxytocin and vasopressin manipulation alter social behavior during the juvenile period in Mongolian gerbils. Developmental Psychobiology, 59(5), 653–657.

Taylor, J. H., & French, J. A. (2015). Oxytocin and vasopressin enhance responsiveness to infant stimuli in adult marmosets. Hormones and Behavior, 75, 154–159.

Taylor, J.H., Walton, J.C., McCann, K.E., Norvelle, A., Liu, Q., Vander Velden, J.W., Borland, J.M., Hart, M., Jin, C., Huhman, K.L. and Cox, D.N., 2022. CRISPR-Cas9 editing of the arginine–vasopressin V1a receptor produces paradoxical changes in social behavior in Syrian hamsters. Proceedings of the National Academy of Sciences, 119(19), pp.e2121037119.

Thompson, E. (2009). The context of female dispersal in Kanyawara chimpanzees. Behaviour, 146(4-5), 629–656.

Trova, S., Bovetti, S., Bonzano, S., De Marchis, S., & Peretto, P. (2021). Sex steroids and the shaping of the peripubertal brain: the sexual-dimorphic set-up of adult neurogenesis. International Journal of Molecular Sciences, 22(15), 7984.

Wang, F., Yin, X. S., Lu, J., Cen, C., & Wang, Y. (2022). Phosphorylation-dependent positive feedback on the oxytocin receptor through the kinase PKD1 contributes to long-term social memory. Science Signaling, 15(719), eabd0033.

Wang, Z., Young, L. J., & Insel, T. R. (1999). Voles and vasopressin: a review of molecular, cellular, and behavioral studies of pair bonding and paternal behaviors. Progress in Brain Research, 119, 483–499.

Wood, R. I., Knoll, A. T., & Levitt, P. (2015). Social housing conditions and oxytocin and vasopressin receptors contribute to ethanol-conditioned social preference in female mice. Physiology & Behavior, 151, 469–477.

Williams, J. R., Catania, K. C., & Carter, C. S. (1992). Development of partner preferences in female prairie voles (Microtus ochrogaster): the role of social and sexual experience. Hormones and Behavior, 26(3), 339–349.

Williams, A.V., Duque-Wilckens, N., Ramos-Maciel, S., Campi, K.L., Bhela, S.K., Xu, C.K., Jackson, K., Chini, B., Pesavento, P.A. and Trainor, B.C. (2020). Social approach and social vigilance are differentially regulated by oxytocin receptors in the nucleus accumbens. Neuropsychopharmacology, 45(9), 1423–1430.

Winslow, J. T., Hastings, N., Carter, C. S., Harbaugh, C. R., & Insel, T. R. (1993). A role for central vasopressin in pair bonding in monogamous prairie voles. Nature, 365(6446), 545–548.

Yohn, C. N., Leithead, A. B., & Becker, E. A. (2017). Increased vasopressin expression in the BNST accompanies paternally induced territoriality in male and female California mouse offspring. Hormones and Behavior, 93, 9–17.

Yohn, C. N., Leithead, A. B., Ford, J., Gill, A., & Becker, E. A. (2018). Paternal care impacts oxytocin expression in California mouse offspring and basal testosterone in female, but not male pups. Frontiers in Behavioral Neuroscience, 12, 181.

Young, L. J., Wang, Z., Donaldson, R., & Rissman, E. F. (1998). Estrogen receptor α is essential for induction of oxytocin receptor by estrogen. Neuroreport, 9(5), 933–936.

Young, L. J., Winslow, J. T., Wang, Z., Gingrich, B., Guo, Q., Matzuk, M. M., & Insel, T. R. (1997). Gene targeting approaches to neuroendocrinology: oxytocin, maternal behavior, and affiliation. Hormones and Behavior, 31(3), 221–231.

Zhao, X., Castelli, F. R., Wang, R., Auger, A. P., & Marler, C. A. (2020). Testosterone-related behavioral and neural mechanisms associated with location preferences: A model for territorial establishment. Hormones and Behavior, 121, 104709.

Zhao, X., & Marler, C. A. (2014). Pair bonding prevents reinforcing effects of testosterone in male California mice in an unfamiliar environment. Proceedings of the Royal Society B: Biological Sciences, 281(1788), 20140985.

Zhao, X., & Marler, C. A. (2016). Social and physical environments as a source of individual variation in the rewarding effects of testosterone in male California mice (Peromyscus californicus). Hormones and Behavior, 85, 30–35.

Zhuang, Q., Zheng, X., Becker, B., Lei, W., Xu, X., & Kendrick, K. M. (2021). Intranasal vasopressin like oxytocin increases social attention by influencing top-down control, but additionally enhances bottom-up control. Psychoneuroendocrinology, 133, 105412.

