## Supplemental Table 1 for "Intranasal oxytocin enhances social preference for parents over peers in male but not female peri-adolescent California mice (*Peromyscus californicus*)"

| **Behavior** | **Type of test behavior was measured in** | **Description** |
| --- | --- | --- |
| *Time in chamber* | Parent-peer preference test (Group 1) | Total number of seconds that a mouse spends in each of the three chambers: left side, right side, and center chamber. Must have all four paws in the side chambers (otherwise still considered time in center chamber). |
| *Time in open arms* | Elevate plus maze (Group 2) | Total number of seconds that a mouse spends with at least two paws on the open arms of the elevated plus maze apparatus. |
| *Crosses through center* | Elevate plus maze (Group 2) | Total number of crosses that a mouse makes through the center of the elevated plus maze apparatus. May be going from dark to light, dark to dark, light to dark or light to light adjacent arms. |
| *Latency to approach* | Novel object (Group 2) | Amount of time that it takes the mouse to first touch the novel object (wire mesh cage). |
| *Investigation time* | Novel object (Group 2) | Total amount of time that the mouse spends in physical contact with the novel object (wire mesh cage). |

**S. Table 1. Ethogram with description of behaviors measured in each test.**
